# E-cigarette vapour from base components propylene glycol and vegetable glycerine inhibits the inflammatory response in macrophages and epithelial cells

**DOI:** 10.1101/2022.03.11.483808

**Authors:** Rachael L. Bell, Danny F. McAuley, Murali Shyamsundar, Cecila M. O’Kane, Yvonne Dombrowski

## Abstract

E-cigarettes are a highly popular nicotine replacement therapy in the process of smoking cessation. Despite this, research on the effects of E-vapours to human health remains limited. The popularity of vaping and mass production of cheap E-liquids has led to compromised safety regulations, with contaminants such as heavy metals and alkaloids detected in multiple liquids. Vaporised E-liquids increase cellular ROS generation and inflammatory cytokine release from pulmonary macrophages. This suggests that E-cigarette usage might activate inflammasomes. Common food additives vegetable glycerine (VG) and propylene glycol (PG) form the base of all E-liquids, but little is known about their inflammatory potential once inhaled. Here, the effect of base components PG and VG on inflammasome activation and cytokine release was investigated in macrophages and epithelial cells exposed to E-liquids and vaporised E-liquid extract (E-vapour). Base E-liquid and E-vapour did not induce cellular cytotoxicity and non-vapourised E-liquid had no effect on IL-8 release. However, basic PG/VG E-vapour inhibited both IL-8 release and conventional inflammasome activation by known inflammatory activators in macrophages and epithelial cells. These results propose a novel inhibitory effect of basic E-vapour components to inflammatory challenges.

## Introduction

Approximately 3.6 million people in the UK use an E-cigarette, alone or in combination, with tobacco smoking, with around two thirds of UK E-cigarette users being ex-smokers (ash.org.uk, 2021). When first appearing on the market, the effects of these devices on the health of individuals was largely unknown. To date it is still unconfirmed if E-cigarettes are a safe alternative to tobacco smoking or another addictive habit that will result in short- and long-term damage to the body. In 2017, E-cigarette sales in San Francisco, California, were banned due to this uncertainty over risk to human health (Article190, 2017). The UK and the EU have not banned the sale of E-cigarettes yet. More research is required into the basic emissions given off by the use of these devices and their effects on human heath in the short and long-term. With a solid understanding of the health implication of these devices, users can make informed decisions on what is the best option for their lifestyle or needs

The basic components (‘base’) of all E-liquids are vegetable glycerine (VG) and propylene glycol (PG) in varying ratios (Hahn., 2014). To this, manufacturers and users can add a variety of flavourings along with varying concentrations of nicotine. Some E-liquids and E-vapours contain carcinogenic chemicals such as aldehydes and nicotine-specific nitrosamines (Khlystov & Samburova, 2016; Klager., 2017; Sleiman., 2016.). There are limited regulations around the manufacture and sale of E-liquids, which are further undermined by the sale and generation of non-regulated E-liquids by users themselves. Effective regulations regarding E-cigarette usage can only be made on the basis on scientific and clinical evidence, highlighting the need for investigation into their effects on human health. This became evident in the summer of 2019 when the new illness termed EVALI, E-cigarette, or Vaping, product use Associated Lung Injury (Siegel., 2019) affected over 2,000 E-cigarette users in America, killing 52 across 26 states (CDC, 2019). The suspected cause was the E-liquid additive vitamin E acetate, a common vitamin supplement safe for topical skin use but not when vaped (Layden., 2019).

The need for more thorough E-cigarette research is especially relevant in context of the COVID-19 pandemic. Indeed, a common flavouring found in E-liquids increased individuals’ susceptibility to SARS-COV-2 infection (Langel., 2022). Additionally, E-vapour increased the expression of ACE2 protein in airway epithelia, which has been linked to the development of COVID-19 (McAlinden., 2021). Nicotine itself increases ACE2 expression (Maggie., 2021) and indeed cigarette smoke has also emerged as a risk factor for COVID-19 development due to increased ACE2 expression. Furthermore, a study which collected peripheral blood mononuclear cells from ‘healthy’ E-cigarette users found increased levels of key proteases linked to ACE2 expression and SARS-COV-2 infection (Kelesidis., 2022).

*In vitro* studies have shown that E-liquids and E-vapours are not as benign as originally believed. E-liquids and -vapours induce cellular toxicity (Farsalinos., 2013 & 2014, Willershausen., 2014, Scheffler., 2015, Hwang., 2016, Leig., 2016, Rowell., 2017, Ween., 2019), inflammatory cytokine release (Wu., 2014, Cervellati., 2014, Rubenstein., 2015, Lerner., 2015, Higham., 2016, Leigh., 2016, Khachatoorian., 2021) and impaired cellular function (Schweitzer., 2015, Hwang., 2016, Scott., 2018, Gómez., 2020) of airway epithelia and immune cells. Human alveolar macrophages secreted increased pro-inflammatory cytokines IL-8, IL-6, and TNFα along with reactive oxygen species (ROS) and were more susceptible to cell death when exposed to E-cigarette vapour extracts (Scott., 2018).

Cell death and ROS are activators of inflammasomes, which regulates the activity and release of pro-inflammatory IL-1β and IL-18 (Martinon, Burns and Tschopp, 2002; Srinivasula., 2002). Inflammasomes are activated upon sensing two distinct danger signals, which determine the type of inflammasome activated. A pathogen associated molecular pattern such as LPS via TLR4 signalling (Creagh and O’Neill, 2006) leads to transcription of inflammasome components such as the Nod-like receptor NLRP3 and apoptosis associated speck like protein (ASC) adaptor along with the pre-cursors of proinflammatory IL-1β and IL-18 (‘priming step’). A second danger signal induces complexing of NLR receptors with inactive capase-1 via the ASC adaptor protein, enabling auto-activation of caspase-1 (Martinon, Burns and Tschopp, 2002; Tschopp, Martinon and Burns, 2003; Mariathasan., 2004). Caspase-1 processes pro-IL-1β and pro-IL-18 along with pro-GSDMD, which forms pores in the cell membrane, allowing the exit of the cytokines and resulting in pyroptotic cell death (He., 2015; Shi., 2015; Chen., 2016; Sborgi., 2016).

E-cigarette users had more inflammasome ASC adaptor protein in their bronchoalveolar lavage (BAL) fluid than never smokers suggesting that inflammasomes might play a role in the inflammatory response to E-cigarettes (Tsai., 2019). Conversely, cinnamaldehyde, a common E-liquid flavouring, reduced the expression of the NLRP3 inflammasome in mice exposed to endotoxin (Xu *et al*., 2017). Despite this, a direct link between E-cigarette use and inflammasome activation in the lung has not been shown

One issue with assessing the safety of E-cigarette vapour on health is the vast variety of over 10,000 E-liquids available. However, all E-liquids contain the base components propylene glycol (PG) and vegetable glycerine (VG). To our knowledge no studies have specifically investigated how these base components PG and VG, that are present in all E-liquids and E-vapours, affect inflammasome activity in cells of pulmonary origin. Here we demonstrate that E-vapour consistent of only the base components PG and VG, have a novel inhibitory role in both cytokine release from epithelial cells and during the priming and activation of macrophage inflammasomes.

## Materials and Methods

### E-liquids and Cigarettes

#### Devices and Liquids

SMOK® Smok Vape Pen Plus 3000mah and Smok VAPE PEN 0.3 ohm Dual Coils 5pcs were purchased from SMOKTECH (Shenzen, China). VG and PG ‘DIY’ Mixing E-Liquid Bases and 72mg/ml nicotine in VG were purchased from Wizard Vapes (Wizard Vapes, London, UK). Reference 3R4F cigarettes were purchased from the University of Kentucky (Kentucky, US)

#### Extract preparation

Atomiser cotton was soaked in 2 ml of ‘basic’ E-liquid (50% PG and 50%VG) in a 50 ml falcon tube for at least 24 hours to avoid dry burning. A new atomiser was used for each E-liquid preparation. The E-cigarette tank and mouthpiece were washed thoroughly with ddH_2_0 after each use and air dried. To create extracts, 50 ml of vapour or cigarette smoke were drawn over 10 ml of cell culture media inside a 50 ml syringe via a silicon tube. The syringe was detached from the tubing, shaken and the vapour/smoke was expelled. This was repeated 30 times for the E-cigarette device, or for the duration of one cigarette. The resulting extract was filtered through a 0.25 μm pore sterile filter in class 2 microbiology safety cabinet and used immediately for cell culture stimulation.

### Cell culture

#### A549 alveolar epithelial-like cells

A549 epithelial cells (Sigma, Darmstadt, Germany) were cultured in Gibco® DMEM (Dulbecco’s Modified Eagle Medium) high glucose, GlutaMAX™ Supplement media (cat. no. 61965026, Thermo Fischer, Massachusetts, US) supplemented with 10% fetal bovine serum (FBS) and 1% Penicillin-Streptomycin (Thermo Fischer, Massachusetts, US). Cells were grown in Nunc™ 75cm^2^ culture flask and maintained at 70-80% confluence. For experiments, cells were plated at 5×10^4^ cells/well in 24 well plates (Thermo Fischer, Massachusetts, US). Cells grew to confluence by 48 hours. The cell culture medium was then changed to 2.5% FBS on the day of the experiment. To induce IL-8 release, cells were treated with TNFα at 10 ng/ml (Peprotech, London, UK) for 24 hours.

#### THP1-ASC-GFP Cells

ASC reporter macrophages (Invivogen, San Diego, US. Cat. Code thp-ascgfp) were cultured in RPMI x1640, supplemented with 10 mM HEPES, 4.5 g/L Glucose, 1 mM Sodium Pyruvate, 10% FBS (Thermo Fischer, Massachusetts, US) and 100 μg/ml Normocin (Invivogen, California, US). Cells were grown in Nunc™ 75cm^2^ culture flasks at a maximum of 2×10^6^ cells/ml with 100 μg/ml of selective antibiotic Zeocin (Invivogen, San Diego, US) added at alternative passages. For experiments, cells were plated at 0.5×10^5^ cells/well in a clear bottom, black walled 96 well plate or at 3×10^5^ cells/well in clear, plastic 24 well plates. Monocytes were differentiated to macrophages with 100 ng/ml of phorbol 12-myristate 13-acetate (PMA) for 72 hours before resting for a further 24 hours without PMA. Cells were washed with phosphate buffered saline (PBS) and stimulated in 2.5% FBS supplemented RMPI.

#### Primary human monocytes

Cells were isolated from human blood using density gradient centrifugation agent Ficoll®-Paque Premium (17-5442-02, GE Healthcare, SIGMA, Darmstadt, Germany). Monocytes were plated at 3×10^5^ cells per well in plastic 24 well plates. Cells were differentiated to macrophages in RPMI x1640 supplemented with 10% FBS containing 10 ng/ml of recombinant human GM-CSF (Peprotech, New Jersey, US) for 7 days. Macrophages were rested for at least 24 hours in fresh RPMI without GM-CSF before being used for experiments.

#### Inflammasome activation

To activate the NLRP3 inflammasome, cells were treated with 1 μg/ml of LPS (Merck, cat.no. L439) for 3 hours. Cells were then washed with dPBS and treated with 20 μM of known inflammasome activator nigericin (Invivogen, California, US) for 2-24 hours.

### Analysis

#### Cytokine detection by ELISA

Human IL-1 beta/IL-1F2 DuoSet ELISA (cat. no. DY201) and Human IL-8/CXCL8 DuoSet ELISA (cat. no. DY208) both R&D systems Bio-Techne Ltd Abingdon, UK) were carried out according to manufacturer’s instructions. Nunc MaxiSorp™ 96 well plates (Thermo Fischer) were read on an Omega Fluostar plate reader (BMG Labtech; Ortenberg, Germany) and analysed using Prism 9 (GraphPad, La Jolla, USA).

#### Western blot analysis of supernatants

Supernatants were concentrated x10 using Amicon Ultra centrifugal units (Merck Millipore, Darmstadt, Germany). The protein content of lysates was analysed using a BCA protein assay kit (Life Technologies; Paisley, UK) according to manufacturer’s instructions. Supernatants and lysates were loaded with sample reducing agent and LDS sample buffer (Invitrogen, Novex, Fischer scientific, UK) in 10 μl, loaded onto a 12% acrylamide gel and separated proteins transferred onto a PVDF membrane (Bio-Rad, Hercules, California, US). Membranes were washed in 1X TBS, 1% Tween (TBST) and blocked in 1X TBST containing 5% Milk powder for 1 hour at room temperature. Anti-caspase-1 (Caspase-1 Polyclonal Antibody, NBP1-45433, Novus biologicals, Bio-Techne Ltd Abingdon, UK) was added according to manufacturer’s recommendations in 5-10 ml of 1X TBST 5% milk at 4°C overnight. Membranes were incubated with HRP-conjugated goat anti-rabbit IgG, diluted to 1:5,000 in 1X TBST 5% milk powder for 1 hour before visualisation with Clarity™ Western ECL Substrate (Bio-Rad, Hercules, California, US) using a Syngene G: BOX with GeneSys software.

#### Cell death analysis

Cytotoxicity Detection LDH Kit (Roche, Indianapolis, USA) was used on cell supernatants according to manufacturer’s instructions. Absorbance values of each well were read on an Omega Fluostar plate reader (BMG Labtech; Ortenberg, Germany) at 405 nm.

#### Live cell imaging of ASC speck formation

THP1-ASC-GFP reporter cells were imaged in black walled, 96 well plates with 100 μl of cell culture medium, using an EVOS™ FL Auto 2 Imaging System (Thermo Fischer, Massachusetts, US). Cells were imaged before and after stimulation. Images were taken at the 2-hour time point using the automatic scan function, taking 4 fields of view per well. ASC specks and green-fluorescent cells were counted manually using ImageJ software (Schindelin., 2012).

#### Statistical Analysis

Data are expressed as mean with standard error of mean. Normal distribution was assessed by KS normality test. Parametric results were interpreted using one-way ANOVA (with Holm-Šídák’s multiple comparisons test) or Student’s *t*-test. Nonparametric results were analysed using Kruskal Wallis H (with Dunn’s multiple comparisons test) or Wilcoxon signed-rank test. Differences of <0.05 were considered significant. All statistical test were performed with GraphPad Prism 9.

## Results

### Basic E-Liquid and basic E-vapour are not cytotoxic to epithelial cells and macrophages

E-liquids were shown to be cytotoxic to pulmonary cells *in vitro* (Scott., 2018). Given the vast number of E-liquid variations it is important to understand what E-liquid components are cytotoxic and if there is a difference between liquids and vapours. Specifically, the base components PG and VG that are present in all variations of E-liquids have not been assessed in detail before and it remains unclear if they themselves pose a health risk. In the following experiments ‘E-vapour’ is referring to the vapour generated from the basic E-liquid components PG and VG only, without any further additives (unless otherwise stated).

Cell death was assessed by lactate dehydrogenase (LDH) release from A549 epithelial cells and primary macrophages from healthy donors that were exposed to 0.03% v/v E-liquid or E-vapour containing only the base components VG/PG or VG/PG plus nicotine.

Basic E-liquids and E-vapour with and without added nicotine did not induce LDH release in A549 epithelial cells (Figure 1A, B) or primary macrophages (Figure 1C, D), compared to untreated controls. E-vapour extracts containing nicotine changed the morphology of primary macrophages suggesting activation but did not induce cytotoxicity (Supplementary Figure 1).

**Figure 1.**
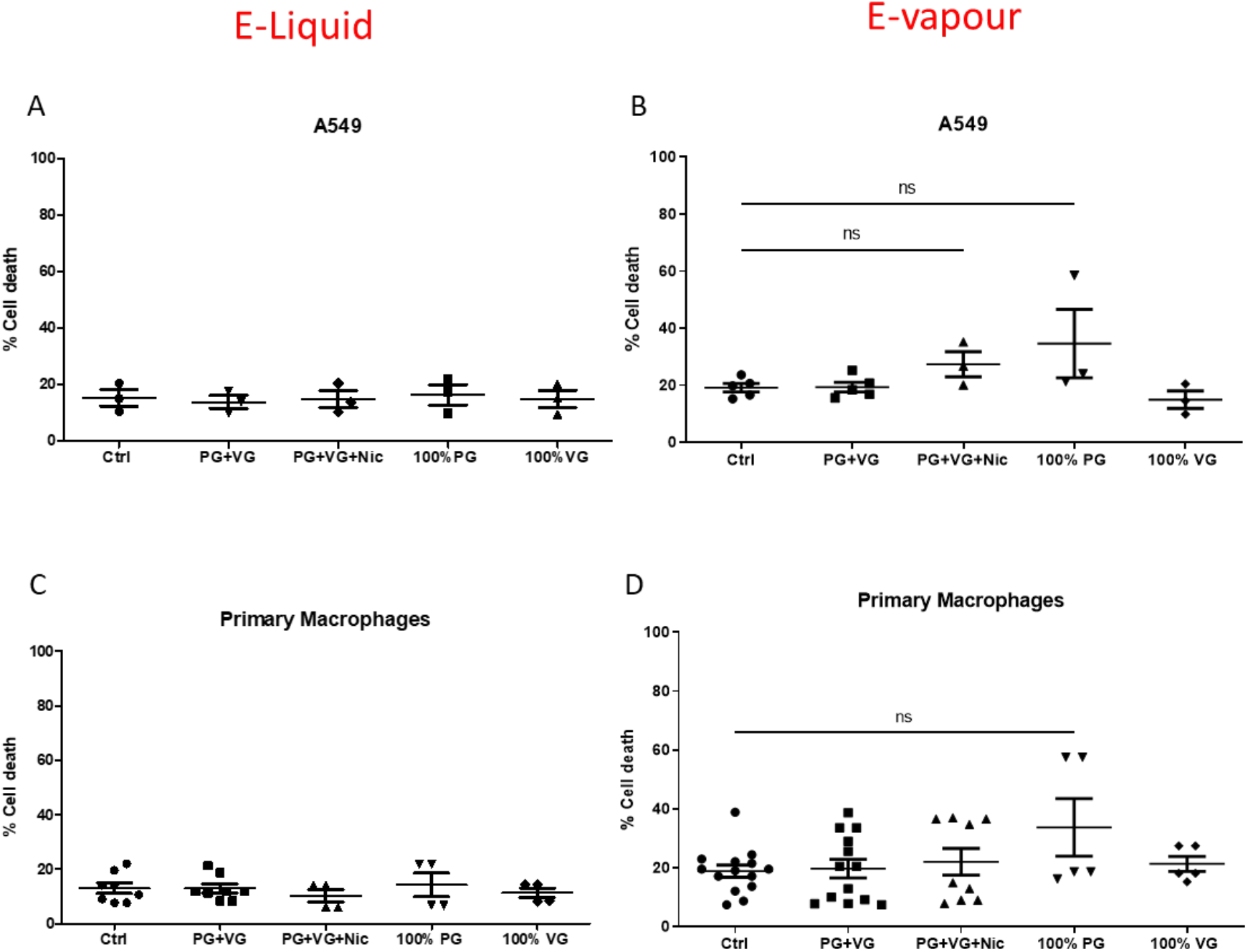
E-Liquid and E-vapour are not cytotoxic to epithelial cells and macrophages. A549 epithelial cells (A, B) and primary macrophages (C, D) were treated for 24 hours with 0.03%v/v E-liquid (A, C) or E-vapour (B, D) and LDH release in the supernatant was assessed. (A,B) n=1-5 independent experiments with 3 technical replicates per experiment; (C,D) n=4-14 healthy donors, with 3 technical replicates per donor. Each data point is an average of 3 replicate values per experiment/donor. Parametric data analysed using 1-way ANOVA (C,D), non-parametric data analysed using Kruskal Wallis H test (A,B). ns: not significant. PG=propylene glycol, VG=vegetable glycerine, nicotine (24mg/ml), 100% PG (0.3% v/v liquid or PG vapour extract), 100% VG (0.3%v/v liquid or VG vapour extract).

### E-vapour inhibits constitutive IL-8 secretion from epithelial cells and macrophages

Lungs of smokers have an increased neutrophil cell count (Schwartz and Weiss, 1994; Higuchi., 2016) likely due to elevated levels of neutrophil chemoattractant IL-8 (Tanino., 2002). Therefore, we investigated if the basic components of E-liquid or E-vapour modulates constitutive IL-8 release from naïve cells.

IL-8 release from A549 lung epithelial cells and primary macrophages was reduced when exposed to E-vapour (Figure 2A, B). Cigarette smoke extract also reduced IL-8 release from primary macrophages to a similar level seen with E-vapour extracts (Figure 2B). However, cigarette smoke extract increased LDH release from these cells indicating increased cell death, which was not seen with E-vapour extracts (Figure 2C). E-liquid did not change constitutive IL-8 release (Figure 2D, 2E).

**Figure 2.**
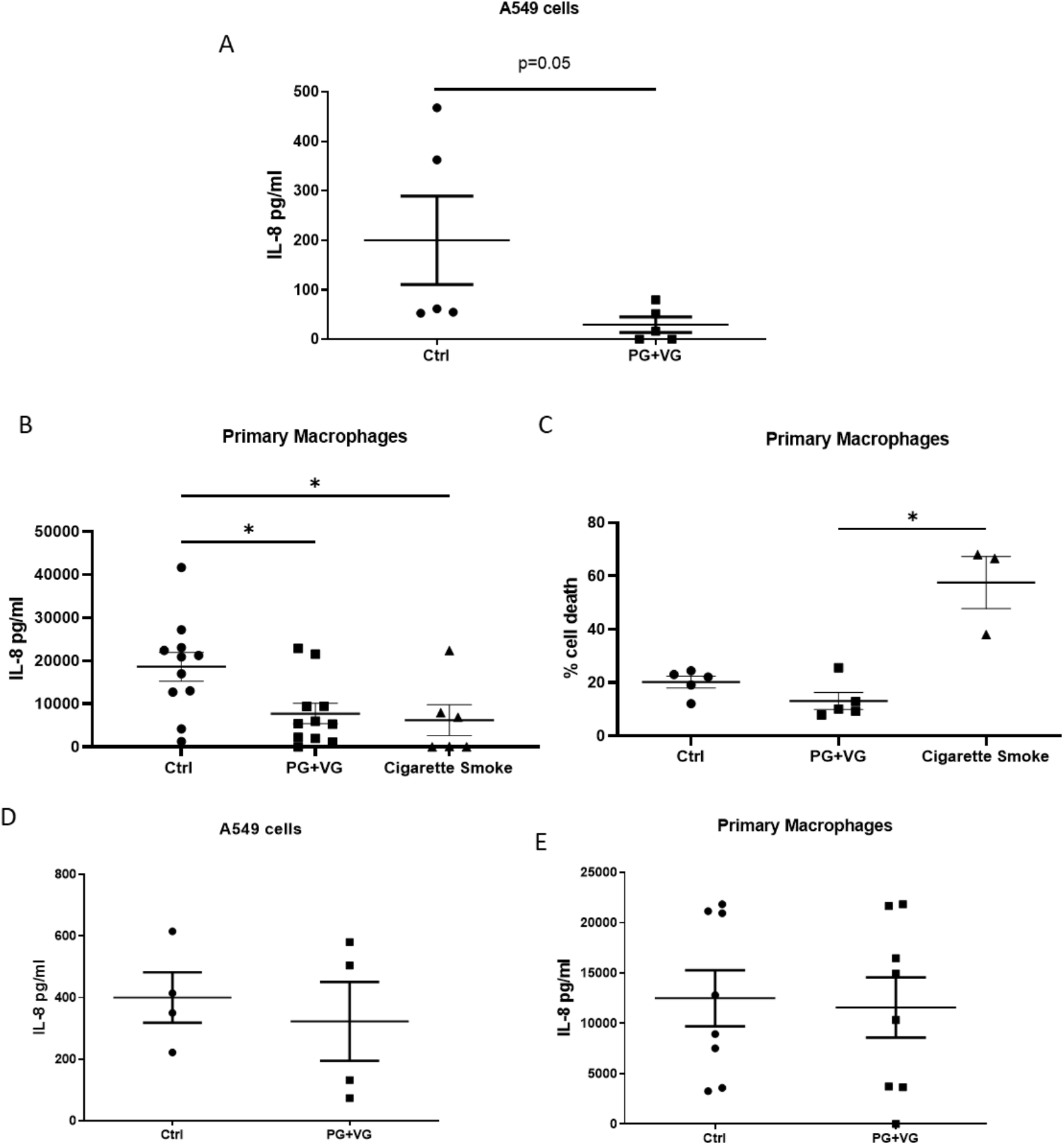
E-vapour inhibits constitutive IL-8 secretion from epithelial cells and macrophages. A549 epithelial cells and macrophages were treated with E-vapour or cigarette smoke extracts (A-C) or with E-liquid (0.03%v/v E-liquid) (D, E) for 24 hours. IL-8 release was analysed by ELISA (A,B,D,E) and cell death was analysed by LDH release (C). (A,D) n=3-5 independent experiments, with 3 technical replicates per experiment, (B, C, E) n=3-11 donors, 3 technical replicates per donor. Each data point is an average of 3 replicate values per experiment/donor *P<0.05. Parametric data (B) analysed using 1-way ANOVA. Non-parametric data analysed using (A) Mann-Whitney U test or (C) Kruskal Wallis H test.

### E-vapour inhibits IL-8 secretion from lung epithelial cells and macrophages after TNFα exposure

To investigate the effects of E-vapour on IL-8 release from activated cells, epithelial cells were stimulated with TNFα in E-vapour extracts. IL-8 release was decreased, although not significantly, when A549 epithelial cells were exposed to TNFα in E-vapour extracts compared to TNFα in control media (Figure 3A).

**Figure 3.**
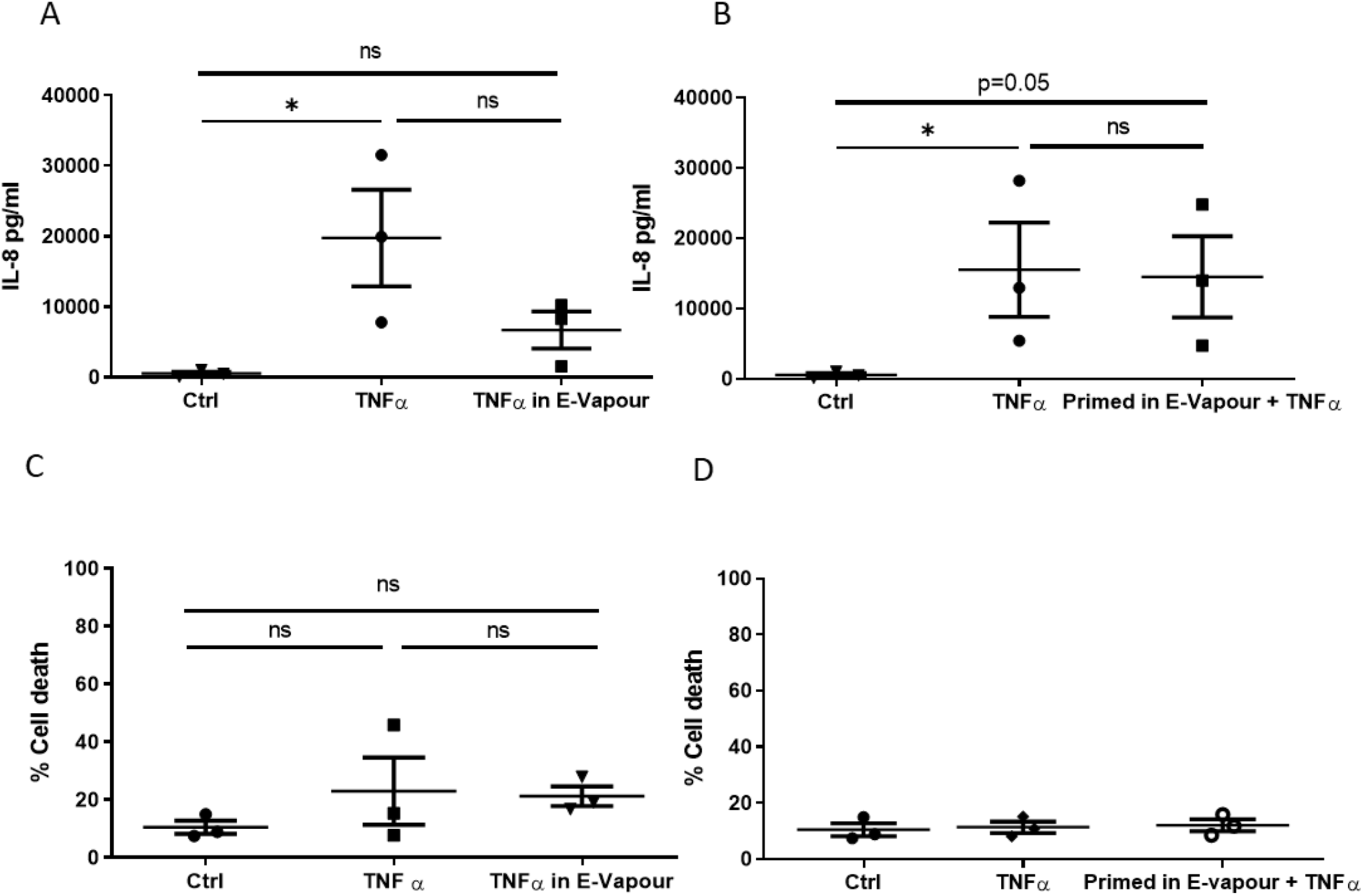
E-vapour inhibits IL-8 secretion from lung epithelial cells and macrophages after TNFα exposure. IL-8 release was analysed by ELISA from A549 epithelial cells treated with TNFα (10 ng/ml, Thermo Fischer) for 24 hours, either simultaneously with E-vapour (A) or after pre-treatment with E-vapour for 2 hours (B). Cell death was analysed by LDH release in cultures simultaneously (C) or consecutively treated with TNFα (D). (A-D) n=3, 3 independent experiments, 3 replicates per experiments, each data point is an average of 3 replicate values per experiment. (A-D) Non-parametric data analysed using Kruskal Wallis H test. *P<0.05, ns: not significant.

To investigate if the reduction in IL-8 release was due to TNFα being blocked by E-vapour components, cells were primed with E-vapour for 2 hours before exposure to TNFα. When primed with E-vapour, there was no difference in IL-8 release with TNFα compared to controls (Figure 3B).

LDH release was assessed to determine if the changes in IL-8 release where due to cell death, however, there was no difference in LDH release (Figure 3 C,D).

The E-vapour mediated reduction in IL-8 release was not due to E-vapour blocking IL-8 detection as spiked recombinant IL-8 was detectable in E-vapour (Supplementary Figure 2A).

### E-vapour inhibits LPS-mediated priming of inflammasomes in macrophages

It is important to understand the effect of E-cigarettes under different physiological conditions. Inflammasomes in immune cells and lung epithelium can be primed and activated by environmental factors inhaled into the airways. Therefore, investigating the impact of E-vapour on cells with already primed and/or active inflammasomes is important to understand the consequences of these devices in different inflammatory states.

To investigate if E-vapours modulate inflammasome priming, THP1 macrophages were primed with LPS for 3 hours with and without E-vapour, then stimulated with nigericin for a further 2 hours to activate the inflammasome (Figure 4A). Cells that were primed with LPS in the presence of E-vapour extract formed fewer ASC specks (Figure 4B, 4D) and released less IL-1β (Figure 4C). E-vapour treated cells had reduced, although not significantly, LDH release indicating less cell death compared to non-E-vapour treated cells (Supplementary Figure 3).

**Figure 4.**
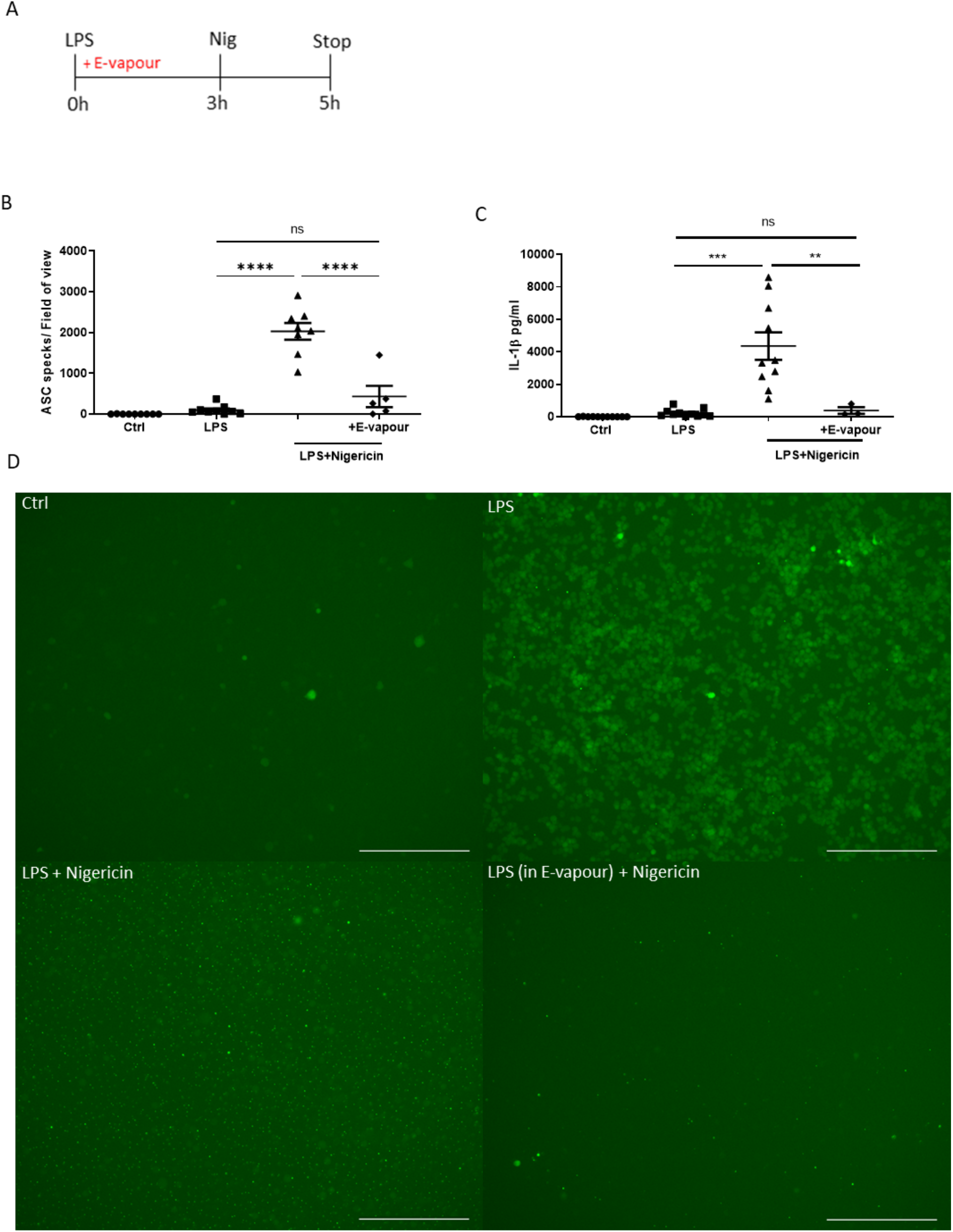
E-vapour inhibits LPS priming of inflammasomes in macrophages. THP1-ASC-GFP reporter macrophages (Invivogen) were primed with LPS (1μg/ml) for 3 hours in the absence or presence of E-vapour, then stimulated for a further 2 hours with nigericin (20μM, Invivogen). (A) Schematic of experimental design. (B) ASC specks were manually counted after cells were imaged on the EVOS™ FL Auto 2 Imaging System and (C) IL-1β release into supernatants was analysed by ELISA. (D) Representative images of B. (B, C) n=3-11 independent experiments, 6 replicates per experiment; each data point is an average of 6 replicate values per experiment. **P<0.005, ***P<0.0005, ****P<0.0001. ns: not significant. (B, C) Parametric data analysed using 1-way ANOVA. All images scale bar =275μm

To determine if this reduced inflammasome activation was a direct or indirect interaction of E-vapour with LPS, E-vapour was added for 3h before LPS priming of cells (Supplementary Figure 4A). Pre-treatment of cells with E-vapour had no effect on ASC specks formed or IL-1β release (Supplementary figure 4B-E).

### E-vapour inhibits nigericin-mediated activation of inflammasomes in macrophages

To investigate the effect of E-vapours on inflammasome activity, LPS primed macrophages were stimulated with inflammasome activator nigericin (Mariathasan., 2006) with or without basic E-vapour (Figure 5A). Macrophages that were stimulated with nigericin in E-vapour had fewer ASC specks (Figure 5B, 5E) and released less IL-1β and IL-18 (Figure 5C, D). E-vapour treated cells had reduced LDH release, although not significant, indicating less cell death compared to non-E-vapour treated cells (Supplementary Figure 5).

**Figure 5.**
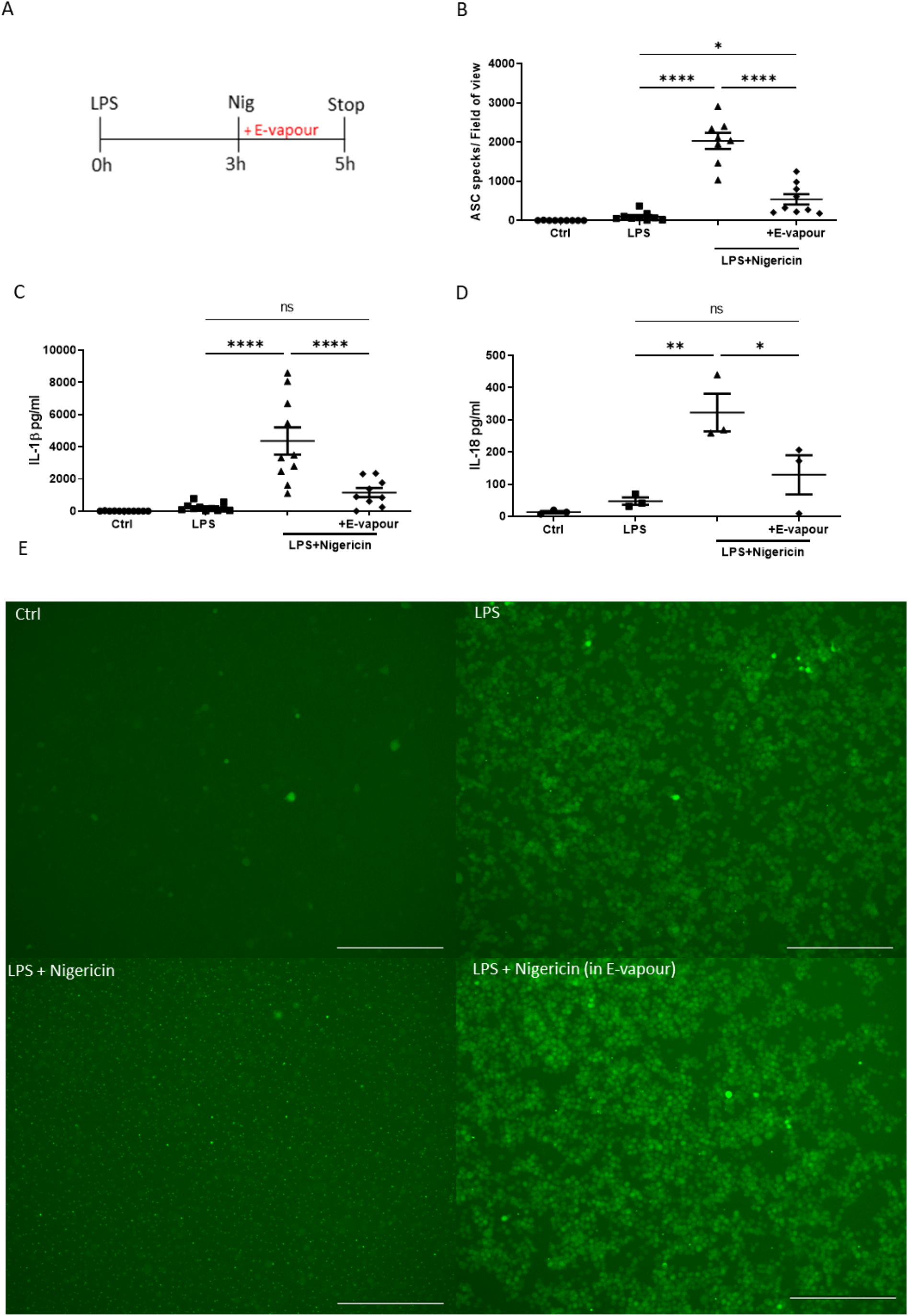
E-vapour inhibits nigericin-mediated activation of inflammasomes in macrophages. THP1-ASC-GFP macrophages (Invivogen) were primed with LPS (1μg/ml, Merck) for 3 hours then stimulated for a further 2 hours with nigericin (20μM, Invivogen) alone or in basic E-vapour. (A) Schematic of experimental design. (B) ASC specks were manually counted after cells were imaged on the EVOS™ FL Auto 2 Imaging System. (C) IL-1β and (D) IL-18 release into supernatants was analysed by ELISA. (E) Representative images of B. n=3-11 independent experiments, 6 replicates per experiment. Each data point is an average of 6 replicate values per experiment. (B-D) Parametric data analysed using 1-way ANOVA. *P<0.05 **P<0.005 ****P<0.0001, ns: not significant. All images scale bar=275μm

To determine if this inhibition was a direct or indirect interaction of E-vapour with nigericin, E-vapour was added for 1.5h before activation of cells with nigericin (Supplementary Figure 6A). Pre-treatment of cells with E-vapour before nigericin activation had no effect on ASC speck formation or IL-1β release (Supplementary figure 6B- E).

### E-vapour inhibits inflammasome activation in primary macrophages

Next, the effects of basic E-vapour on priming and activation of inflammasomes was investigated using primary macrophages from healthy donors. Macrophages were exposed to basic E-vapour as described in the experimental design either in conjunction with LPS (Figure 6A) or in conjunction with nigericin (Figure 6B).

**Figure 6.**
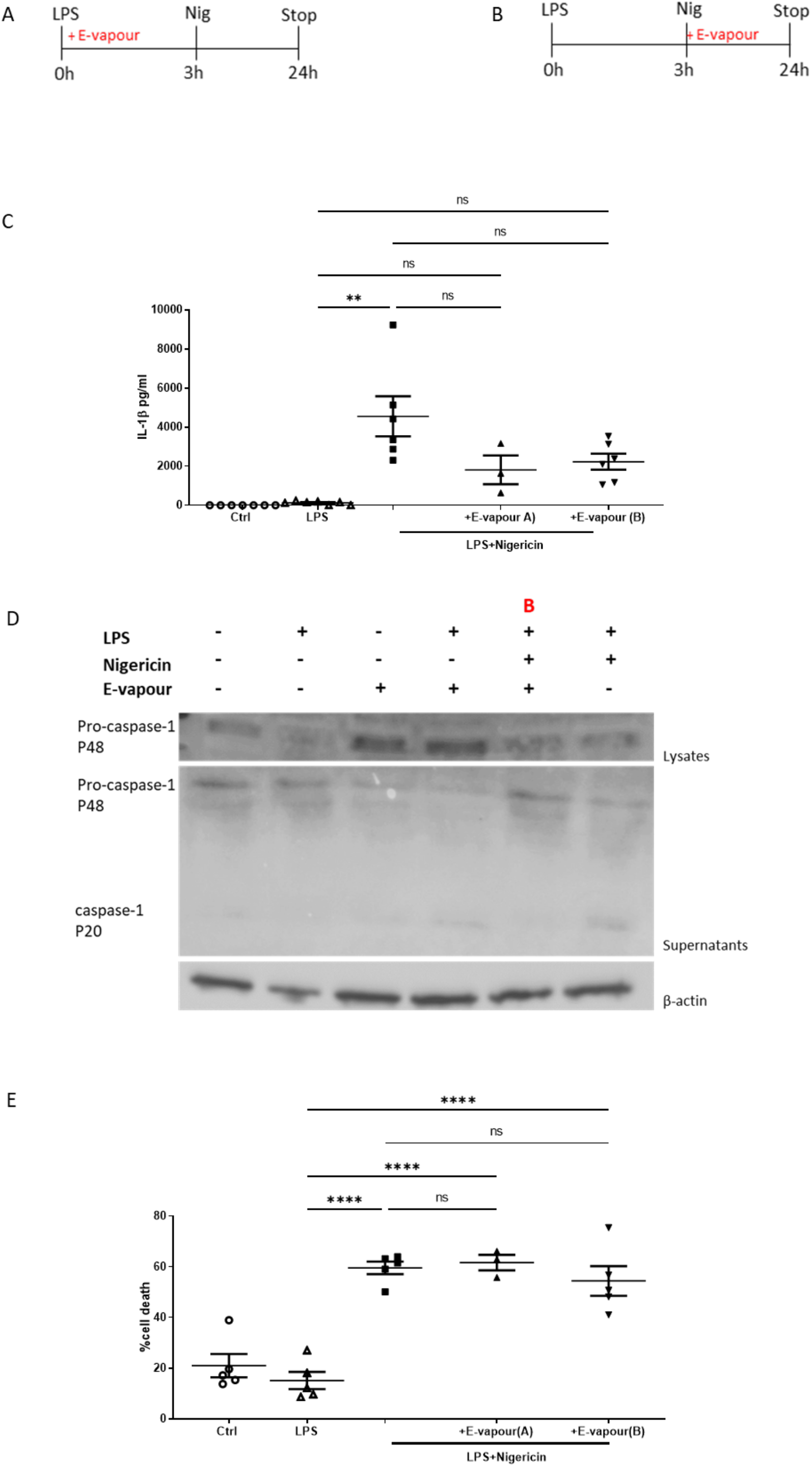
E-vapour reduces inflammasome activation in primary macrophages. Primary monocytes were differentiated with GMCSF (10ng/ml Peprotech) to macrophages for 5 days. Macrophages were primed for 3 hours with LPS (1μg/ml, Merck) and activated with nigericin (20μΜ, Invivogen) for a further 24 hours, with or without E-vapour extract, respectively. (A, B) Schematics of experimental designs. (C) Supernatants were analysed for IL-1β release by ELISA. (D) Caspase-1 was analysed by western blot from lysates (top) and supernatants (middle) of cells treated as described in (B) (indicated by B, β-actin in lysates (bottom) served as loading control). (E) Cell death was analysed by LDH release. n=3-7 donors, 3 replicates per experiment. Each data point is an average of 3 replicate values per experiment. (C, E) Non-parametric data analysed using Kruskal Wallis H test. **P<0.005 ****P<0.0001. ns: not significant.

There was an observed, but not significant, decrease in the amount of IL-1β released from macrophages when E-vapour was present during either the inflammasome priming or activation stage compared to when macrophages were primed and activated with LPS and nigericin on their own (Figure 6C).

Additionally, active caspase-1, which requires inflammasome activity, was absent from the supernatants of macrophages when inflammasomes were activated in the presence of E-vapour (Figure 6 D).

There was no difference in LDH release from macrophages in the presence or absence of E-vapour during inflammasome priming and activation (Figure 6E).

### Emissions from the E-cigarette atomiser dampen inflammasome activation in macrophages

We found that basic E-vapour consistent of only PG and VG had an inhibitory effect on activated cells (Figures 2–6). These effects where not seen with un-vapourised E-liquids (Figure 2–3). When E-liquid is vaporised, it is possible that emissions may be given off by the heating of the metal atomiser and cotton wick. To test if these emissions played a role in the inhibitory effects mediated by basic E-vapour, a new atomiser was fitted to the E-cigarette and PBS was vaporised as a vehicle to carry any emissions from the device itself. THP macrophages were primed with LPS and the inflammasome was activated with nigericin; vapourised PBS was added with either LPS (Figure 7A) or nigericin (Figure 7B).

**Figure 7.**
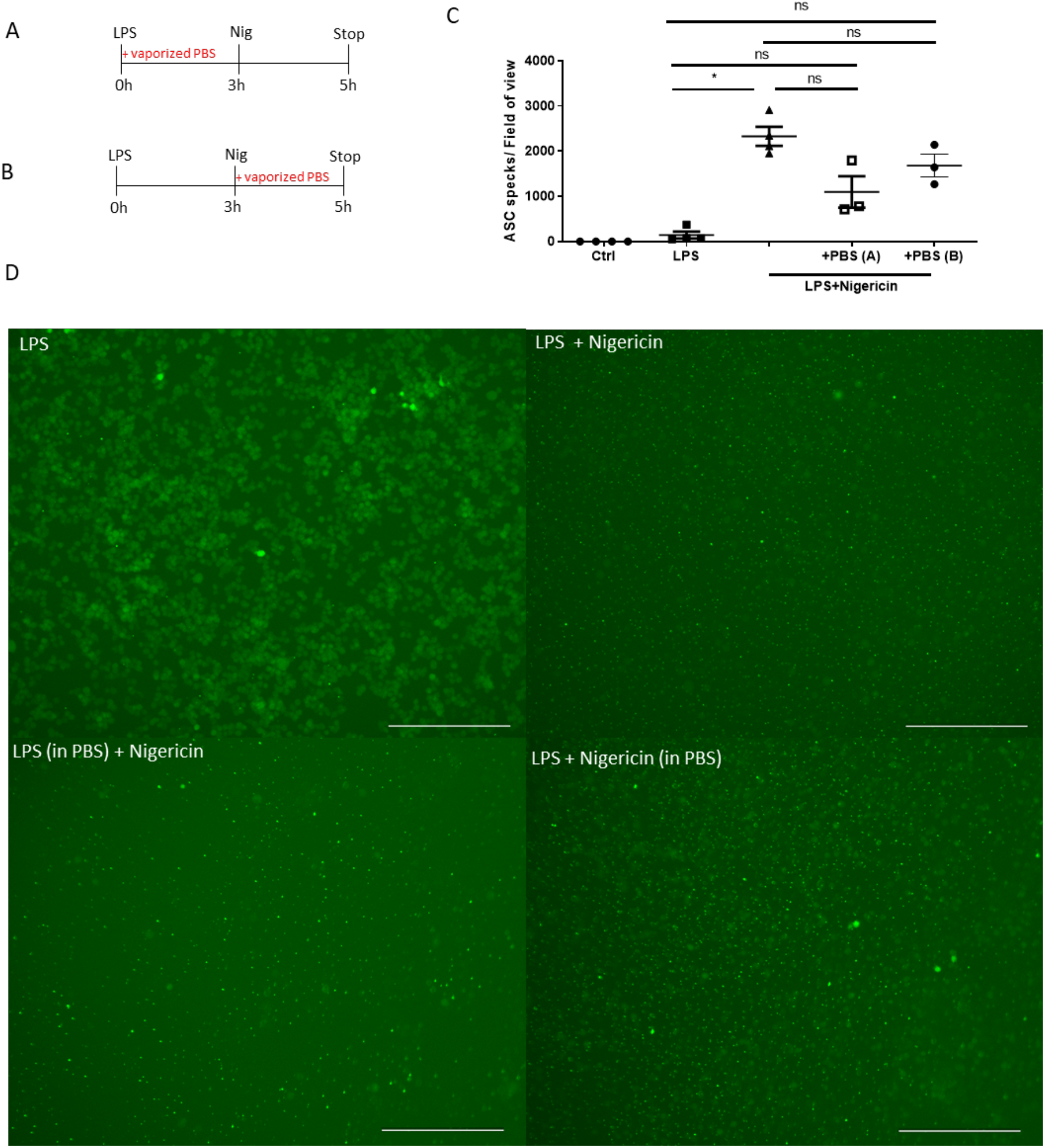
Emissions from the E-cigarette atomiser dampen inflammasome activation in macrophages. THP1-ASC-GFP macrophages (Invivogen) were primed for 3 hours with LPS (1μg/ml, Merck) then activated with nigericin (20μΜ, Invivogen) for a further 24 hours, with and without vapourised PBS extract. (A, B) experimental design. (C) Macrophages were imaged on the EVOS™ FL Auto 2 Imaging System and ASC specks counted (D) representative images of (C). n=3-4 independent experiments, 6 replicates per experiment. Each data point is an average of 6 replicate values per experiment. Non-parametric data analysed using Kruskal Wallis H test. *P<0.05; ns: not significant. All images scale bar=275μm

The addition of vapourised PBS with either LPS or nigericin reduced the amount of ASC specks formed compared to the positive control, however this did not reach statistical significance (Figure 7C,D).

## Discussion

Various E-vapours can induce inflammatory cytokine release such as IL-8, IL-6, TNFα (Cervellati., 2014; Wu., 2014; Rubenstein., 2015; Leigh., 2016; Scott., 2018), cytotoxicity (Farsalinos,., 2013; Willershausen., 2014; Scheffler, 2015; Scott., 2016), ROS generation (Rubenstein., 2015; Scott., 2018) and gene expression alterations in a variety of cells including bronchial cells and alveolar macrophages (Martin., 2016; Lee., 2017). Many studies demonstrating a pro-inflammatory response to E-vapour were nicotine and flavouring-dependant (Bahl., 2012; Lerner., 2015; Leigh., 2016; Yu., 2016; Ghosh., 2019). Yet, it is not known if the basic E-liquid components, PG and VG, that are present in all E-liquids and E-vapour themselves have pro-inflammatory functions. Although PG and VG are deemed safe to ingest as food additives and topically as cosmetics, their use in inhaled E-liquids may have unknown consequences. Additionally, the vaporisation of E-liquid changes its chemical composition (Cheng, 2014; Khlystov., 2016) and increases its cytotoxic effects in alveolar macrophages (Scott et al., 2016).

Neither basic E-vapour containing only PG and VG, or nicotine-containing E-vapour or E-liquid were cytotoxic to epithelial cells or macrophages tested here (Figure 1). However, all E-vapours reduced IL-8 release from epithelial cells and macrophages, both in naïve state and activated with TNFα (Figure 2 and 3). Cigarette smoke also reduced IL-8 release from primary macrophages and led to increased macrophage death, which was not observed with E-vapour (Figure 2). Furthermore, we observed a change in macrophage morphology exposed to nicotine-containing E-vapours compared to macrophages with non-nicotine contain E-vapours (Supplementary figure 1). This implies that although cell death and cytokine release was similar between nicotine and non-nicotine containing cells, nicotine could possibly induce intracellular adverse effects, not detected by secretions into the supernatant of cultures.

Our data indicate that although not cytotoxic, E-vapour interferes with a central inflammatory response such as IL-8 signalling, a key inflammatory cytokine, responsible for activation of immune cells, recruitment of systemic neutrophils and the subsequent induction of inflammatory clearance at the site of injury or infection (Baggiolini, 1993; Nocker., 1996; Dinarello, 2000).

The role of inflammasome activation in the inflammatory response to E-vapour is not well characterised. There is indirect evidence of inflammasome activation by E-vapour stimulation in cell models, by IL-1β release (Leigh., 2016; Ween., 2019) and by ASC presence in the BAL of E-cigarette users (Tsai., 2019), although to our knowledge, inflammasome activation has never been directly studied in relation to E-vapour. We found E-vapour inhibited inflammasome priming by LPS (Figure 4,6) and activation by nigericin in inflammasome reporter cells and primary macrophages (Figure 5,6). E-vapour mediated inhibition of inflammasome priming was dependent on a direct interaction with LPS, as pre-stimulation and removal of E-vapour prior to LPS priming had no effect (Supplementary figure 4). The same was observed with inflammasome activation, where pre-treatment of primed cells with E-vapour prior to signal 2 had no effect on nigericin-mediated activation of the inflammasome (Supplementary Figure 6). The inhibitory effects of E-vapour on inflammasome priming and activation were confirmed in primary macrophages (Figure 6). Additionally, when added simultaneously with nigericin, E-vapour inhibited the release of active caspase-1 into the supernatant, confirming the inhibition of inflammasome activation (Figure 6).

Here we show for the first-time that basic E- vapour consistent of just the base components PG and VG can supress priming and activation of inflammasomes in the presence of a classical inflammasome activators such as LPS or nigericin. This is not the first evidence of E-cigarette ingredients supressing inflammatory responses. Morris *et al* demonstrated significantly decreased IL-1β, TNFα, IL-6 and IL-8 from LPS activated THP-1 macrophages with multiple flavouring compounds commonly used in E-liquids (Morris., 2021). Here, however, we show that the vapourised base ingredients present in all E-liquids can negatively regulate critical inflammatory responses even in the absence of additives, which has potentially significant implications for all E-cigarette users. Gomez *et al* showed a similar phenomenon where THP-1 macrophages infected with *Mycobacterium tuberculosis* exposed to unflavoured nicotine-free E-vapour demonstrated significantly less phagocytic activity and reduced IL-1β and TNFα release (Gomez., 2020). The ability of basic E-vapour to inhibit inflammatory signalling and functionality of activated and infected cells has consequences for pathogen clearance and may result in more systemic, prolonged infections.

Nigericin is a NLRP3 inflammasome inducer and the NLRP3 inflammasome is most likely to be involved in inhaled airway challenges such as E-vapour. While inhibition of inflammasome activation by basic E-vapour has not been reported before, NLRP3 inflammasome inhibition by cigarette smoke has been described (Morris., 2015; Han., 2017; Ye., 2019); NLRP3 protein was reduced via ubiquitin mediated proteasomal processing in THP1 cells and C57BL/6 mice exposed to cigarettes smoke, and the NLRP3 inflammasome was supressed in reaction to *Candida albicans* in a rat model exposed to cigarette smoke. Indeed, Morris *et al* showed the NLRP3 inflammasomes response to asbestos was inhibited by cigarettes smoke, while we show here that nigericin fails to activate primed macrophages in the presence of basic E-vapour.

Many studies asses the safety of E-vapour in comparison to cigarette smoke. Despite similar reductions in IL-8 release, cigarette smoke owes much of this reduction to high cell toxicity, whereas basic E-vapour is not cytotoxic to the cells tested here (Figure 1). It is also important to note the structural differences between smoke and E-vapours. Although particle size of smoke and vapour is similar (Williams., 2013), the amount of particles produced by conventional smoke is greater than vapour (Zhang., 2013) the longevity of the particles in the air differs (Martuzevicius., 2018) and E-cigarette users also tend to use an E-cigarette more frequently (Eissenberg, 2010; Vansickel., 2010), increasing their frequency of exposure to potentially harmful products of vaporisation.

The vaporisation of E-liquid is a key factor for these observed inhibitions in cellular activation and cytokine release, as E-liquid addition did not have the same effect (Figure 2, 3). This may be due to emissions given off by the metallic atomiser as vapourised PBS reduced inflammasome activation in THP1 macrophages, albeit not to the same extent (Figure 7). Two studies showed that the emissions of electronic cigarettes contained traces of metal, which potentially originate from the metallic atomiser of the E-cigarette device (Williams., 2017; Kamilari., 2018). In this context, the suppression of the canonical and non-canonical NLRP3 inflammasome by heavy metals (Ahn ., 2018) could be relevant. It is thus possible that our observed effects are results of different sources of emissions that originate from the heating of elements such as the metallic atomiser.

Our data are highly relevant in the context of the SARS-COV-2 pandemic. E-cigarette vapour increased the virulence of key lung pathogens in epithelia (Gilpin., 2019), decrease antimicrobial activity of neutrophils and epithelia (Hwang., 2016) and increase viral load of pulmonary cells (Wu., 2014). This coupled with decreased inflammatory responses from activated immune cells makes E-cigarette use a potentially important risk factor in the contraction and spread of the SARS-COV-2 virus.

In summary, we show a novel role for E-vapour from basic E-liquid components PG and VG to supress IL-8 production from epithelial cells and macrophages, along with suppression of inflammasome activation in macrophages. These data warrant further investigation into the mechanisms involved in dampening this inflammatory response, especially in the context of bacterial and viral infection, before recommending these devices as a safe adjuvant to smoking cessation.

## Acknowledgements

We would like to thank Dr Deirdre Gilpin for the help with setting up the E-cigarette model. RB was funded by a PhD studentship from the Department for Education, Northern Ireland.

