## supplementary figure 1-6 for "E-cigarette vapour from base components propylene glycol and vegetable glycerine inhibits the inflammatory response in macrophages and epithelial cells"

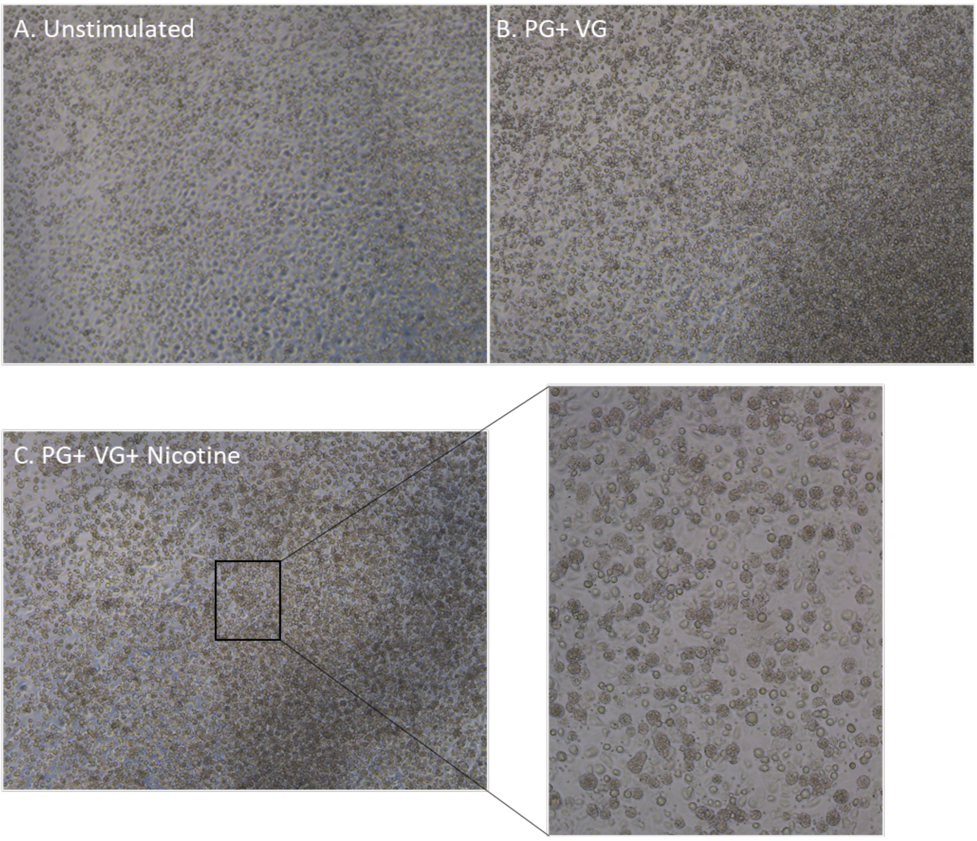


**SFig1: Nicotine changes macrophage morphology**

*Primary macrophages were differentiated with GM-CSF (10ng/ml, Peprotech) for 5 days and treated with basic E-vapour extracts for 24 hours. (A) Untreated macrophages, (B) PG+VG vapour treated macrophages, (C) PG+VG+nictoine treated macrophages.PG=propylene glycol, VG=vegetable glycerine, nicotine (24mg/ml).*

Supplementary figure 2:

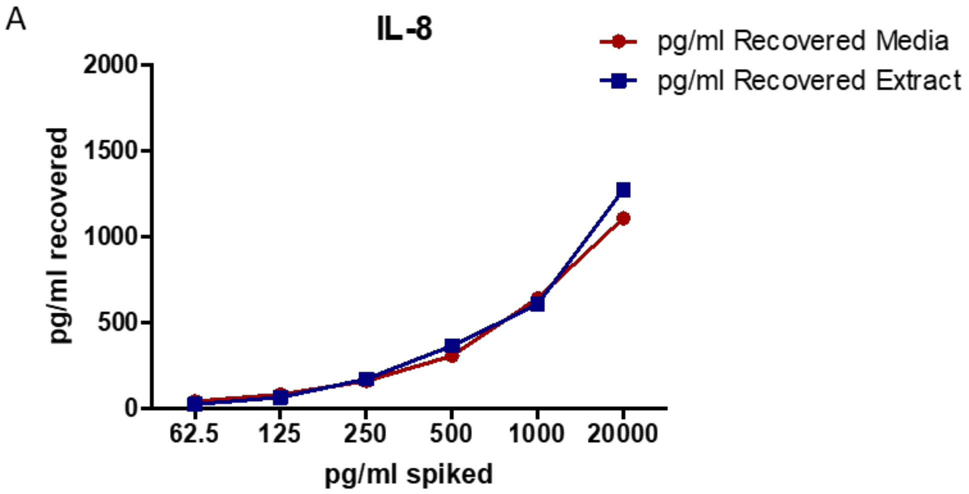


**SFig2: IL-8 detection is not blocked by E-vapour**

*Human recombinant IL-8 (62.5-2,000pg/ml) was spiked into cell culture media or E-vapour extract at 5% $CO_2$, 37$^{o}$C and measured after 24h by ELISA.*

Supplementary Figure 3:

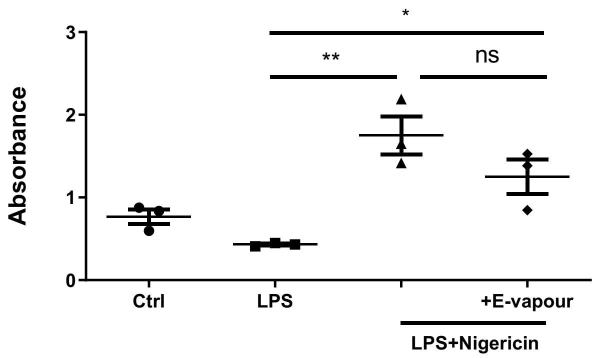


**SFig3: LDH release from inflammasome activated and E-vapour treated macrophages**

*THP1-ASC-GFP reporter macrophages (Invivogen) were primed with LPS (1µg/ml) for 3 hours in the absence or presence of E-vapour, then stimulated for a further 2 hours with nigericin (20µM, Invivogen). LDH release was measured in the supernatant. n=3 independent experiments, 6 replicates per experiment; each data point is an average of 6 replicate values per experiment. Parametric data analysed using 1-way ANOVA *P<0.05 **P<0.005. ns: not significant.*

Supplementary Figure 4:

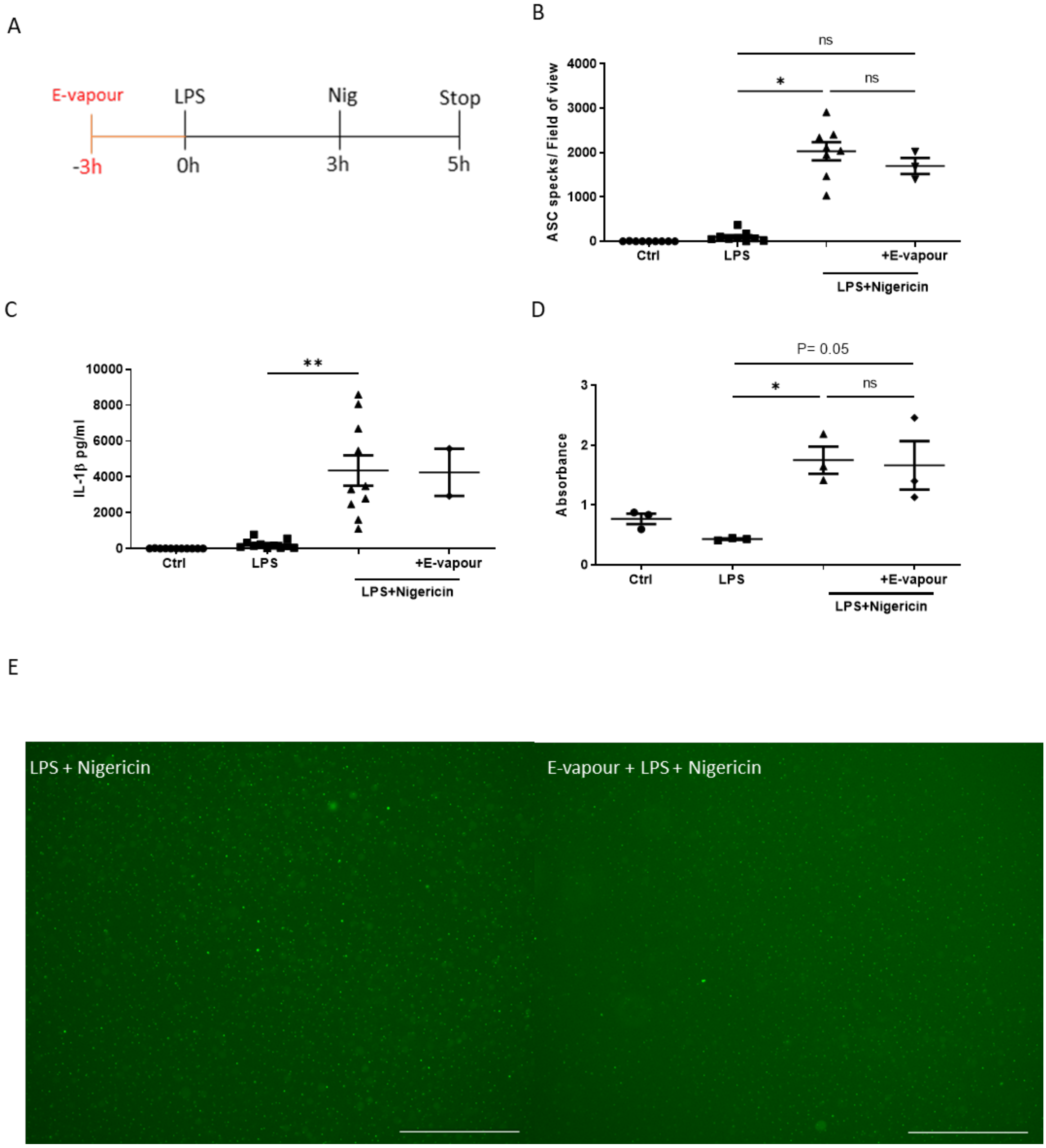


**SFig4. E-vapour pre-treatment does not inhibit inflammasome activation**

*THP1-ASC-GFP reporter macrophages were pre-treated with E-vapour extracts for 3 hours before LPS (1µg/ml) was added for 3 hours then stimulated for a further 2 hours with nigericin (20µM). (A) Experimental design, (B) ASC specks were manually* *counted after cells were imaged on the EVOS™ FL Auto 2 Imaging System, (C) IL-1β release in the supernatant was analysed by ELISA and (D) cell death was analysed by LDH release. (E) Representative images of B-D. (B-D) n=2-11 independent experiments, 6 replicates per experiment. Each data point is an average of 6 replicate values per experiment. (B-D) Non-parametric data analysed using Kruskal Wallis H test. *P<0.05; **P<0.005; ns: not significant. All images scale bar=275μm*

Supplementary Figure 5:

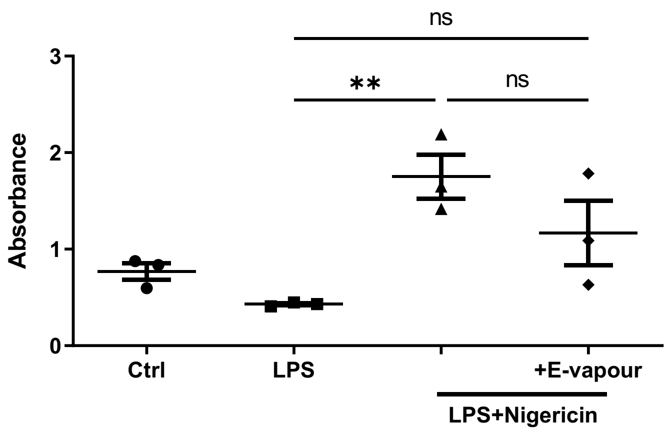


**SFig5: LDH release from inflammasome activated and E-vapour treated macrophages**

*THP1-ASC-GFP macrophages (Invivogen) were primed with LPS (1µg/ml, Merck) for 3 hours then stimulated for a further 2 hours with nigericin (20µM, Invivogen) alone or in basic E-vapour. Cell death was analysed by LDH release. n=3 independent experiments, 6 replicates per experiment. Each data point is an average of 6 replicate values per experiment. Parametric data analysed using 1-way ANOVA. **P<0.005, ns: not significant.*

Supplementary figure 6:

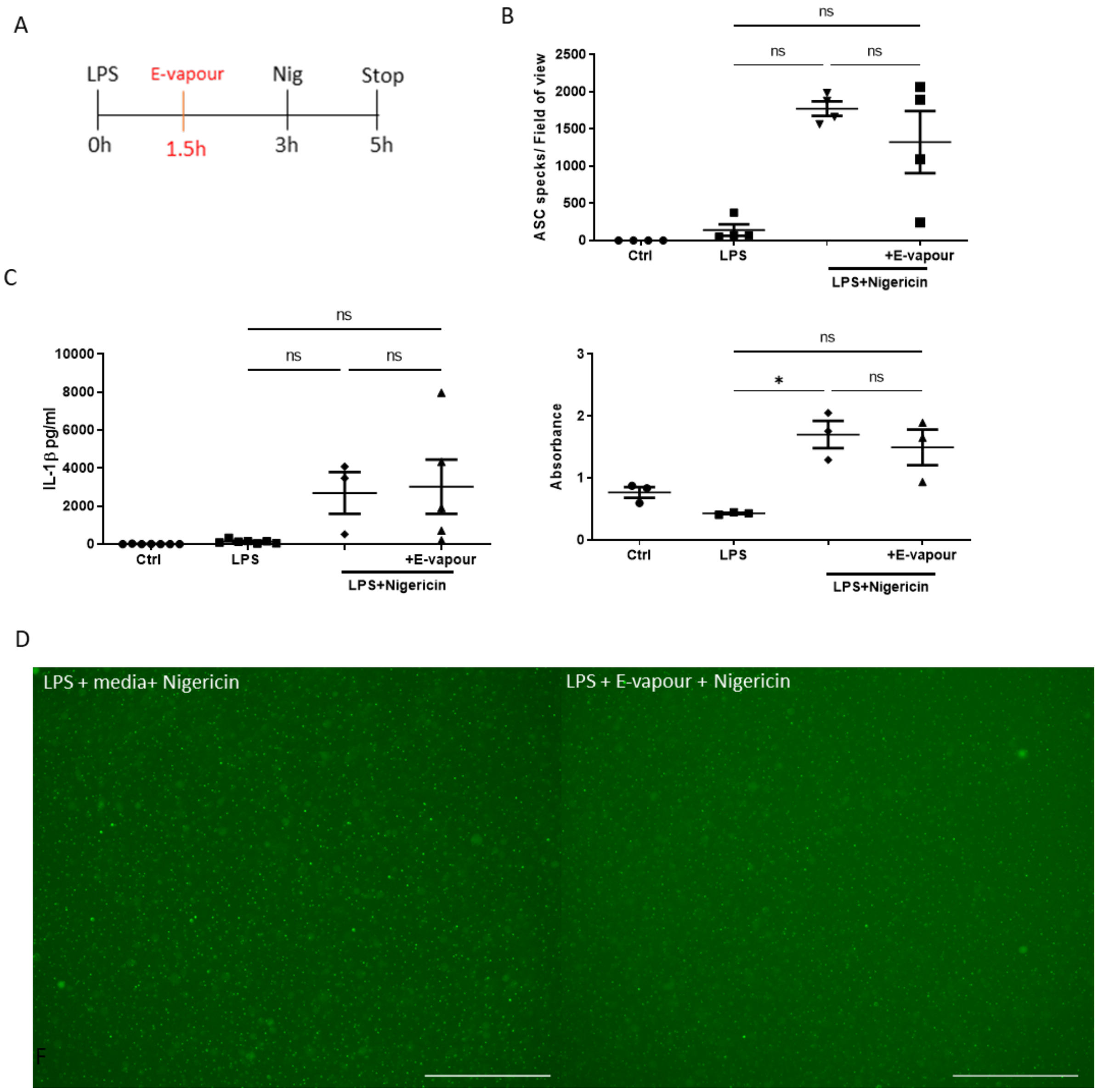


**SFig6. E-vapour pre-treatment does not inhibit inflammasome activation in macrophages**

*THP1-ASC-GFP macrophages (Invivogen) were primed with LPS (1µg/ml, Merck) for 3 hours then stimulated for a further 2 hours with Nigericin (20µM, Invivogen). E-vapour was added 1.5 hours before nigericin stimulation. (A) Schematic of experimental design. (B) ASC specks were counted after cells were imaged on the*

*EVOS™ FL Auto 2 Imaging System. (C) IL-1β was analysed in supernatants by ELISA and (D) cell death was analysed by LDH release. (E) Representative images of B.*

*n=3-7 independent experiments, 6 replicates per experiment, each data point is an average of 6 replicate values per experiment. *P<0.05; ns: not significant. Non-parametric data analysed using Kruskal Wallis H test. All images scale bar=275μm*
